# Depletion of FH, an essential TCA cycle enzyme, drives proliferation in a two-step model

**DOI:** 10.1101/2022.06.16.496410

**Authors:** Balakrishnan Solaimuthu, Michal Lichtenstein, Arata Hayashi, Anees Khatib, Inbar Plaschkes, Yuval Nevo, Mayur Tanna, Shirel Lavi, Ophry Pines, Yoav D. Shaul

## Abstract

Several tumor suppressor genes do not follow the canonical function of cell cycle repressors. For example, fumarate hydratase (FH) is an evolutionary conserved TCA cycle enzyme that reversibly catalyzes the hydration of fumarate to L-malate and has a moonlight function in the DNA damage response (DDR). Interestingly, despite this enzyme’s essential role in central carbon metabolism, FH is inactive or absent in several tumors, such as hereditary leiomyomatosis and renal cell cancer (HLRCC). Accordingly, FH has a contradictory cellular function, as it is pro-survival through its role in the TCA cycle, yet its loss can drive tumorigenesis. These observations have supported the role of FH as a tumor suppressor. Here, we solved this contradiction by determining the molecular mechanisms that allow the cells to survive and even proliferate upon FH loss. We found that the cells’ response to FH loss is separated into two distinct time frames based on cell proliferation and DNA damage repair. During the early stages of FH loss, the cells’ proliferation and DNA damage repair are inhibited. However, over time the cells overcome the FH loss and form knockout clones, indistinguishable from WT cells in their proliferation rate. Due to the FH loss effect on DNA damage repair, we assumed that the recovered cells bear adaptive mutations. Therefore, we applied whole-exome sequencing to identify such mutated genes systematically. Indeed, we identified recurring mutations in genes belonging to the central oncogenic signaling pathways, such as JAK/STAT, which we validated to be impaired in FH-KO clones. Intriguingly, we demonstrated that these adaptive mutations are responsible for FH-KO cell proliferation under TCA cycle malfunction.

## Background

Fumarate hydratase (FH) is a TCA cycle enzyme that catalyzes the hydration of fumarate to L-malate ^1^. Despite its essential role in central carbon metabolism, FH is inactive or absent in several tumors, such as hereditary leiomyomatosis and renal cell cancer (HLRCC) ^2^. This lack of functional FH results in the significant rewiring of the cellular metabolism as the cells accumulate fumarate and demonstrate a fermentative metabolism, which includes reduced oxygen consumption and elevated lactate secretion ^3^. Fumarate accumulation in cells functions as an “oncometabolite” ^4^ which inhibits α-ketoglutarate (αKG)-dependent dioxygenases, histone DNA demethylases ^5^, and stabilizes the hypoxia-induced factor 1 alpha (HIF1α) ^6^. Moreover, fumarate accumulation induces the epithelial-mesenchymal transition (EMT) program ^7^ that is associated with high-grade tumors ^8^ by altering the epigenetic landscape of these cells. Together, FH has contradictory cellular functions, as it is pro-survival through its role in the TCA cycle and functions as a tumor suppressor in tumorigenesis. However, a comprehensive cellular mechanism to explain these opposing roles of FH is still missing.

In addition to its localization in the mitochondria, FH is also present in the cytosol/nucleus, where it participates in the DNA damage response (DDR) ^9^. In the nucleus, FH converts malate to fumarate, which, upon the induction of DNA breaks, inhibits the KDM2B-mediated H3K36me2 demethylation, resulting in non-homologous end-joining repair ^10^. However, the function of fumarate in the DDR is still unclear, as it suppresses the DDR by inhibiting KDM4B ^11^. Here we show that the effect of FH loss in cells is divided into two-time frames. In the early stages of FH loss, the cells are under stress, do not proliferate, and inhibit the DDR. Consequently, this loss causes the accumulation of mutations in the FH-KO cells, enriched within genes encoding components of oncogenic pathways. These adaptive mutations provide the long-term FH-KO cells with the capability to proliferate even in the absence of a functional TCA.

## Results

### Short-term FH loss inhibits cell proliferation

We first sought to determine the impact of FH loss on the cell proliferation rate and in particular characterizing the difference between short-term and long-term effects. We found that FH expression levels in the non-cancerous human embryonic kidney cell line (HEK-293T) cells were almost completely abolished after five days from CRISPR-Cas9 induction and after one day with the hepatocellular carcinoma (HepG2) cells (Figure 1A). Nevertheless, after a few days of the CRISPR-Cas9 induction, we observed that in the polyclonal population of FH-KO cells, the enzyme expression gradually returned to WT levels. These findings are consistent with the role of FH in proliferation, as cells that have escaped the knockout and still express FH, have a growth advantage over those lacking the enzyme (FH-KO). This conclusion is further supported by the short-term knockout of the DPYD, a non-essential metabolic gene ^8^, which even after seven days retained a low expression level (Figure S1A), since there is no selection for its function as for FH. Importantly, despite this impact on cell proliferation, we could generate stable FH-KO clones (FH-Clone) for both cell lines (Figure 1A), suggesting a long-term cellular mechanism that enables cells to overcome FH loss.

**Figure 1:**
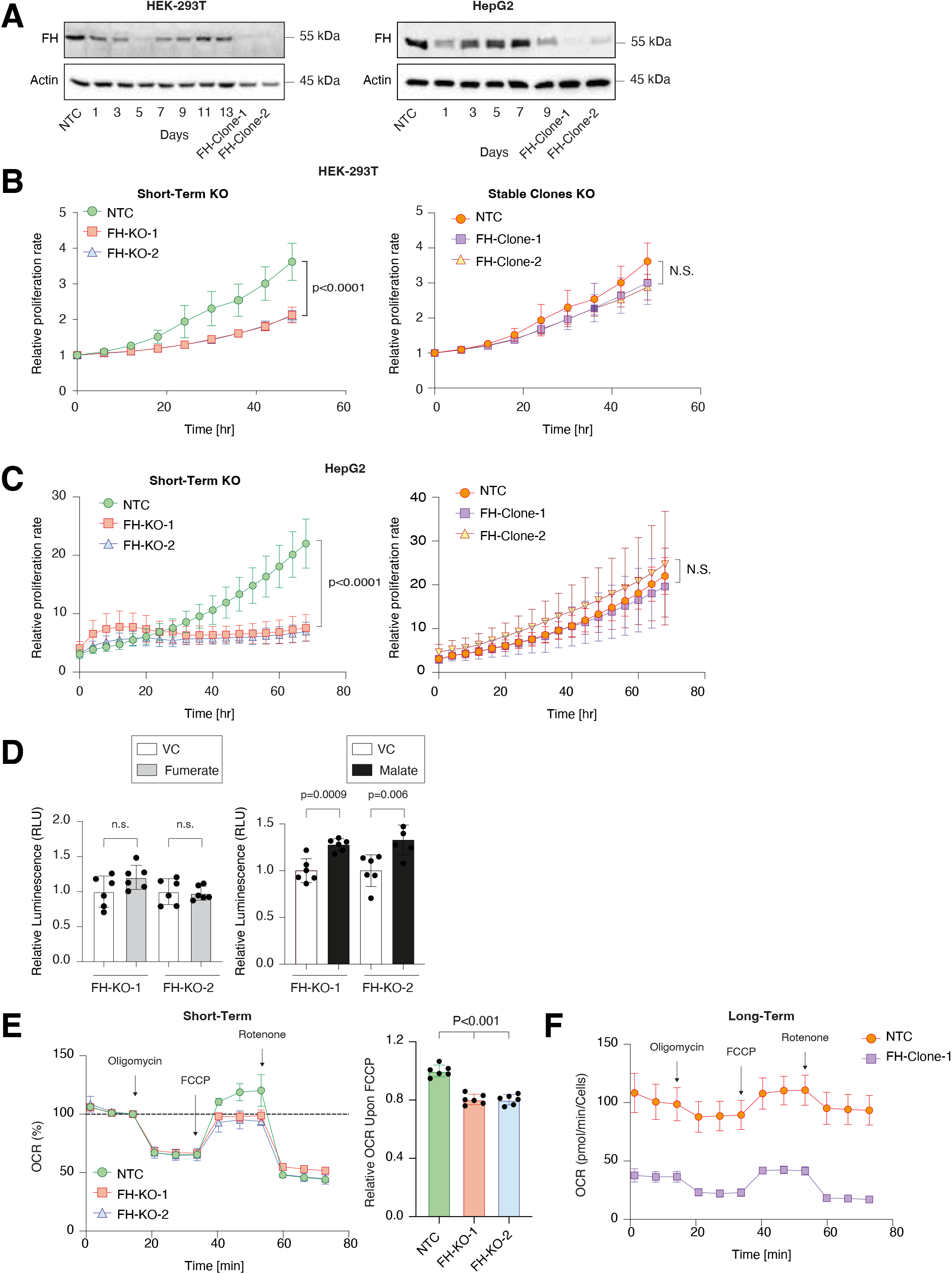
Short-term FH loss inhibits cell proliferation. (**A**) FH expression profile in short-term and long-term knockout cell lines. FH was silenced in both HEK-293T and HepG2 using the CRISPR-Cas9 system and then subjected to 24 hours of puromycin selection. At the indicated times post-selection (short-term), the cells were lysed and subjected to immunoblotting using the indicated antibodies. FH-Clone-1 and FH-Clone-2 are long-term FH knockout clones and NTC is a non-targeted control. (**B**) FH was knocked out from HEK-293T, and then two days post puromycin selection, the cells were subjected to IncuCyte for proliferation analysis and monitored every 2 hours up to 48 hours (left). To determine the long-term effect, two stable FH-KO clones were subjected to the same procedure for 48 hours (right). NTC-is non-targeted control. The analysis of both short-term and long-term was conducted simultaneously. Each bar represents the mean ± SD for n=6. The p-value was determined by Student’s *t*-test. N.S.= no statistically significant differences between the samples. (**C**) HepG2 cells were analyzed similarly to HEK-293T above by IncuCyte for 70 hours. Each bar represents the mean ± SD for n=6. The p-value was determined by Student’s *t*-test. N.S.= no statistically significant differences between the samples. (**D**) The proliferation rate of FH short-term knockouts was not rescued by supplementing fumarate (mono-ethyl fumarate) and only partially rescued by malate (diethyl malate). FH was knocked out in HEK-293T cells and then treated with vehicle control (DMSO, VC), fumarate [250 µM], or malate [250 µM] for three days. The p*-*values were determined by Student’s *t*-test. (**E**) The relative oxygen consumption rate (OCR) of NTC and two different short-term FH-KO HepG2 cells was determined using Seahorse. Atthe indicated time the cells were treated with the different drugs (arrows). The dash bar indicates the baseline of NTC cells (left). Quantification of the relative OCR upon FCCP treatment is represented as a graph (right). Each bar represents the mean ± SD for n=6. The p-value was determined by Student’s *t*-test. (**F**) OCR levels of NTC HepG2 and long-term FH-KO clones.

We conducted real-time monitoring of the effect of FH loss on the cell growth using the Incucyte Live-Cell analysis system. We found that in both cell lines, the proliferation rate of the two different short-term FH-KO cells was significantly lower than WT cells (Figures 1B, C, and S1B). Moreover, supplementing the medium with fumarate (FH substrate) had no effect (Figure S1C), while malate (FH product) supported FH-KO cell growth (Figure 1D). Therefore, the initial response of cells to FH loss, is inhibition of proliferation due to TCA cycle impairment. Nevertheless, several short-term FH-KO cells can overcome this proliferation obstacle and convert into long-term growing clones.

Next, we determined the effect of FH loss on cell metabolism. We found that the oxygen consumption rate (OCR) basal respiration level was similar in both HepG2 WT and short-term FH knockout cells (Figure 1E), but is significantly reduced in long-term clones (Figure 1F). Importantly, the short-term FH loss affected the OCR, as it significantly reduced the maximal respiration levels (Figure 1E). Since measuring the OCR in HEK-293T is challenging, we restricted our experiments to HepG2. Collectively, we reveal that the impact of FH loss on cell proliferation is divided into two phases, based on the time from CRISPR-Cas9 induction; short-and long-term. Furthermore, the ability of cells to proliferate without a functional FH, demonstrates that these cells have adapted to the metabolic change.

### Short-term FH loss induces DNA damage

Recent studies indicated that FH cellular function is more complex than initially thought, as it is also localized in the nucleus, where it functions in the DDR ^1,9^. Indeed, the immediate effect of FH KO (short-term) in both HEK-293T (Figure 2A) and HepG2 (Figure 2B) was stepwise induction in H2A.X phosphorylation (γ-H2AX), which is negatively correlated with FH loss. In contrast, this phosphorylation, a sensitive marker of DNA double-strand breaks^12^, is absent in the long-term FH-KO cells, suggesting that DNA damage is impaired at the early stages of FH loss. This supports our notion that DNA damage functions as an integral part of the cell adaptation to FH loss.

**Figure 2:**
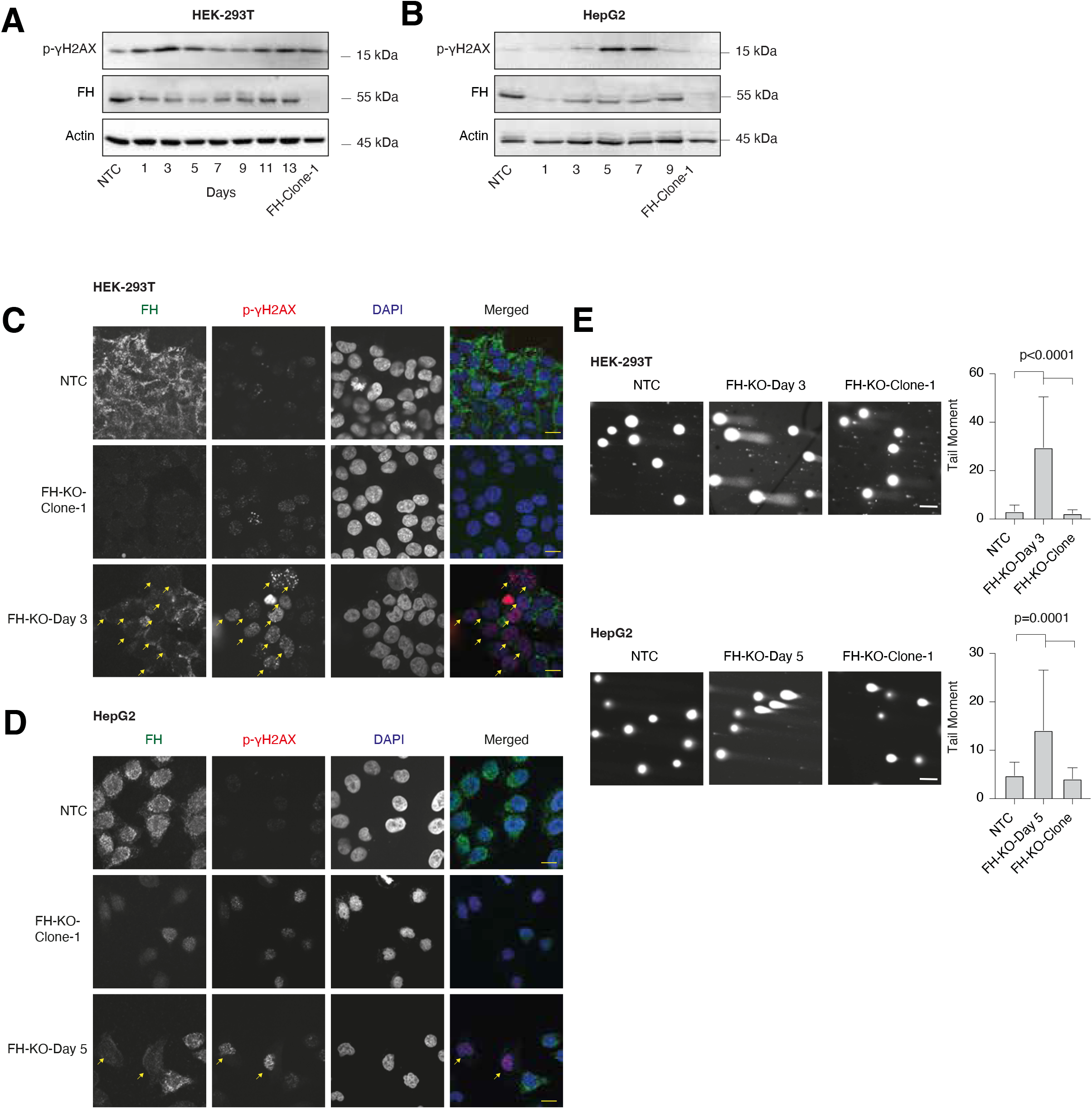
Short-term FH loss induces DNA damage: (**A**) γH2AX phosphorylation is elevated in short-term FH loss in HEK-293T cells. FH expression was silenced in HEK-293T using the CRISPR-Cas9 system. On each of the indicated days (post-infection) the cells were lysed and subject to Western blot using the indicated Abs. FH-KO-1 are stable long-term FH knockout clones. (**B**) γH2AX phosphorylation is elevated in short-term FH depleted HepG2 cell line as analyzed in A. (**C**) HEK-293T cells were fixed and stained for FH (green), p-γH2AX (red), and DAPI (blue). Bar=10µm. Arrows indicate FH knockout cells. (**D**) HepG2 cells analyzed as in C. (**E**) Short-term FH loss induces DNA breaks. NTC, short-term, and long-term FH-KO cells in both HEK-293T and HepG2 were subjected to the comet assay. Representative images of each sample (left). Bar=10µm. Quantification of data is reported as the number of tail moments (right); each bar represents the mean ± SD for n≥30. The p-value was determined by Student’s *t*-test.

Co-staining of HEK-293T and HepG2 cells with FH and γ-H2A.X antibodies indicates that the DNA damage is specific for the short-term KO cells (Figure 2C and D). Fascinatingly, in this polyclonal population (short-term), only cells that appear as FH-KO (not stained with FH antibodies) exhibit γ-H2A.X foci. In line with these results, we show by comet assay that the short-term FH-KO cells demonstrate a significantly high level of DNA breaks versus WT and long-term clones (Figure 2E). Since the long-term FH-KO cells do not display DNA damage, despite the absence of functional FH, this suggests that the growth of these cells is a product of an adaption process. Hence, identifying the mutations induced by the FH-KO-dependent DNA damage impairment will help understand the adaption process.

### Whole exome sequencing analysis of long-term FH KO clones

To assess the consequence of the impaired DNA damage response, we subjected 11 HEK-293T FH-KO clones and one WT (as a reference) to whole-exome sequencing. The mutations identified in the different clones were compared to WT cells using Mutect2, a somatic variant caller ^13^ (Figure 3A, illustration). This analysis resulted in 14,959 FH-KO-specific variants. Next, we filtered out the variants localized in intronic regions using the wANNOVAR online annotation tool ^14^. To avoid the possibility of non-specific variants, we first retained only those with more than ten reads. Then, to increase the probability of true hits, we focused only on the variants that exhibited a high percentage of reads from the total. To this end, for every variant in each FH-KO clone, we calculated the percentage of alteration out of the total reads of this sequence. This analysis allowed us to identify those with at least 30% variation reads, about 12% of the total variants (Figure S2).

**Figure 3:**
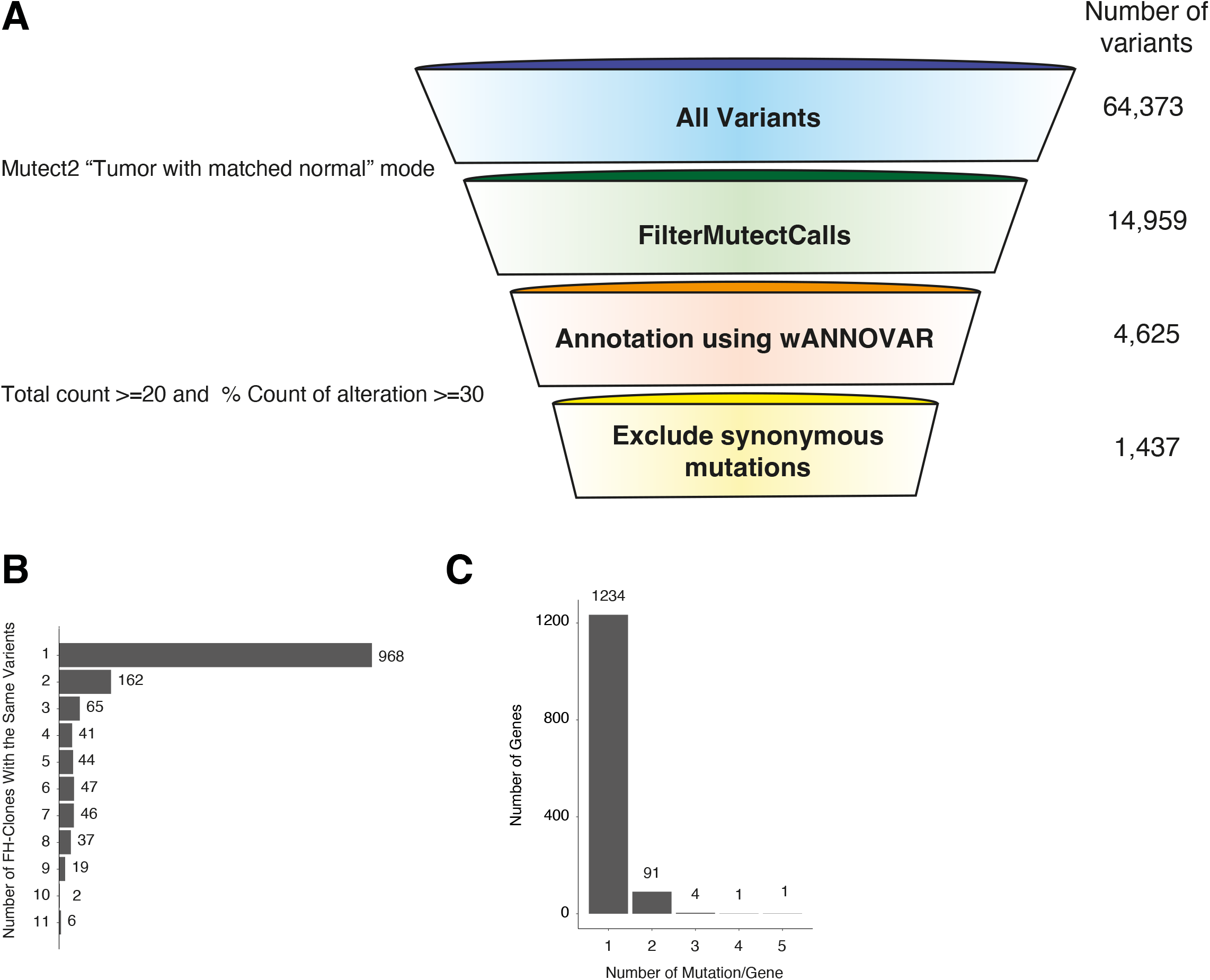
Exome sequencing analysis of FH knockout cells: (**A**) Whole-genome exome sequencing analysis and variant filtering. Variant filtering funnel scheme with the number of variants left after each filtering (right). (**B**) A column bar graph representing the number of FH-clones with the same variants. The number on the right of the bar indicates the number of the variants. (**C**) A column bar graph representing the number of mutations in each gene. The number on the top of the bar indicates the number of genes.

Next, we removed all synonymous mutations, resulting in 1,437 variants. We found that the majority of the genes in the list had one mutation per clone (968 variants) (Figure 3B). However, several genes had recurrent mutations in multiple clones, including six mutated in all samples (MTRF1L, IGFALS, CTSD, BMP1, PCDHB13, and FAM196A). Moreover, 97 genes were mutated in two or more locations (Figure 3C). Interestingly, we observed that the mutated genes did not distribute randomly over the chromosomes, as several are significantly more abundant on chromosomes 9, 11, and 19 (Figure S3). These results portray a novel perception of how FH loss induces genomic mutations, which presumably provide the cells with the ability to overcome the FH-related proliferation impairment.

### Long-term FH knockout clones harbor mutations in central metabolic and signaling pathways

We inquired whether mutations detected in the previous section are associated with common biological processes. Using the Kyoto encyclopedia of genes and genomes (KEGG) pathway database (https://www.genome.jp/kegg/pathway.html), we discovered that these FH loss-associated mutations are present in genes of multiple metabolic pathways, such as central carbon metabolism, nucleotide biosynthesis, and amino acid metabolism (Figure 4A). Moreover, we identified that FH loss significantly induces mutations in signaling pathways (Figure S3A and S3B), indicating that they provide a growth advantage. Specifically, these pathways include central pro-survival signaling cascades such as the MAPK, chemokine, Wnt, and JAK/STAT signaling (Figure 4B). By applying the KEGG mapper color tool, we present the mutated genes in MAPK-signaling, chemokines, and JAK-STAT pathways (Figure 4C). These findings highlight the enrichment of FH-loss-induced mutations in central signaling pathways, indicating that these genomic impairments are not random.

**Figure 4:**
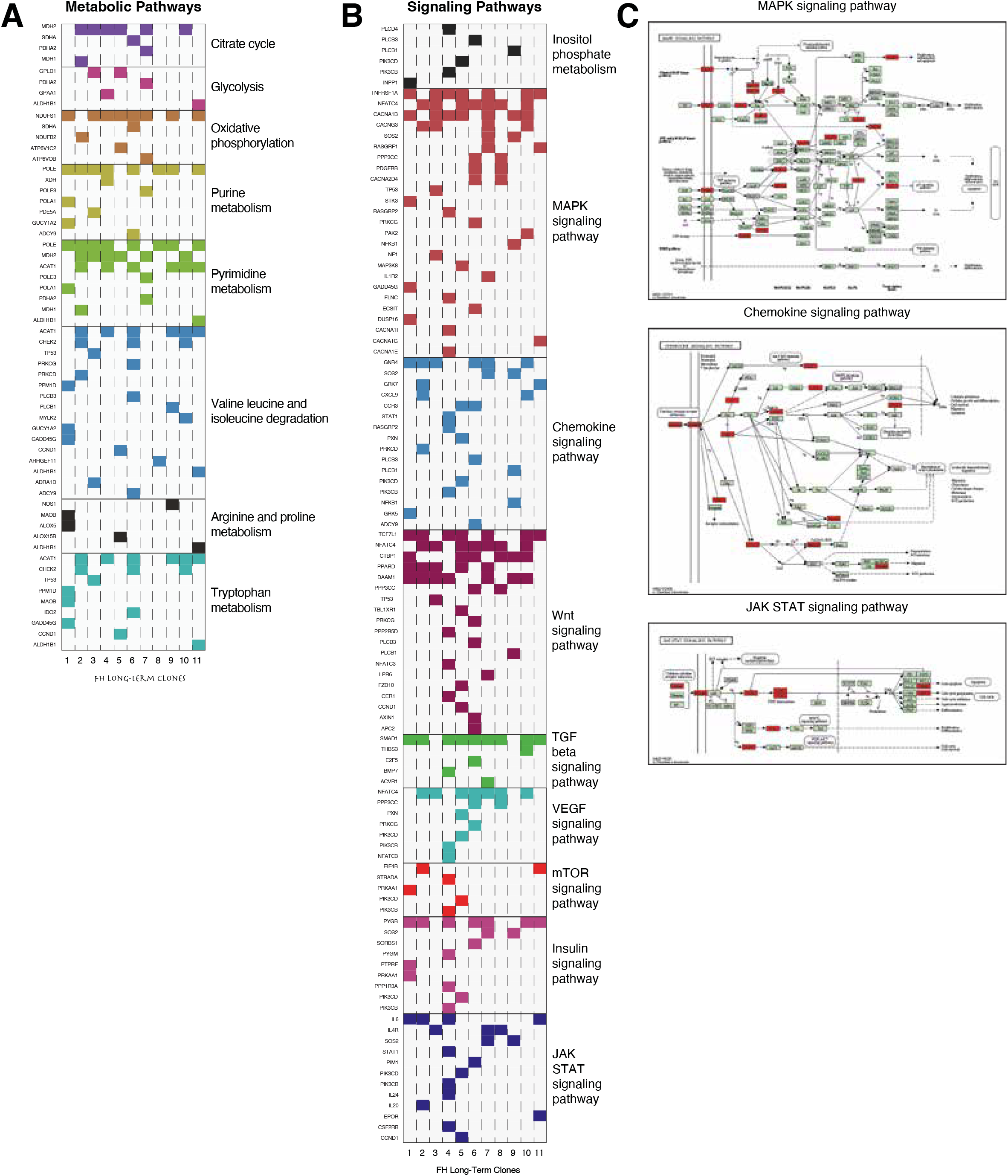
Long-term FH loss is accompanied by mutations in selected signaling and metabolic genes: (**A**) The mutated genes are presented as metabolic pathways. Each column represents a different FH-KO clone and each row represents a different gene. The different pathways are color-coded and were determined by the KEGG database. (**B**) The names of selected signaling pathways are color labeled as in A. (**C**) KEGG map displays the distribution of the mutated genes (marked in red) in the different signaling pathways.

### FH loss results in STAT3 signaling impairment

To further examine the effect of FH loss on signaling pathways, we chose the well-known JAK/STAT3 which may provide a step in the adaption process. We demonstrated that this signaling pathway regulates HEK-293T growth, since expressing a constitutively activated STAT3 (A662C, N664C, V667L, CA-STAT3) ^15,16^, significantly inhibited its proliferation rate (Figure 5A). Accordingly, FH-loss inhibited STAT3 activity, as reflected by tyrosine 705 phosphorylation (Y705) reduction (Figure 5B), and repression of its downstream target, IL-6 (Figure 5C). Finally, RNA seq analysis of the WT and FH-KO clone followed by Gene Set Enrichment Analyses (GSEA)^17^, demonstrated that FH loss significantly reduces the “Hallmark of IL6-JAK-STAT3 signaling”. Additionally, we noticed other STAT3-related hallmarks such as “Hallmark of the epithelial-mesenchymal transition”, and “Hallmark of KRAS signaling-up”, were reduced (Figure 5D). Interestingly, analysis of patient data using the Cbiopotal web-based (https://www.cbioportal.org) tool, identified two tumor samples; MPCPROJECT_0130 (Prostate Adenocarcinoma), and SD5038 (Cutaneous Melanoma), in which both IL-6 and FH are deleted. Collectively, we demonstrated that in FH-KO cells, the STAT3 signaling is impaired, which restored proliferation.

**Figure 5:**
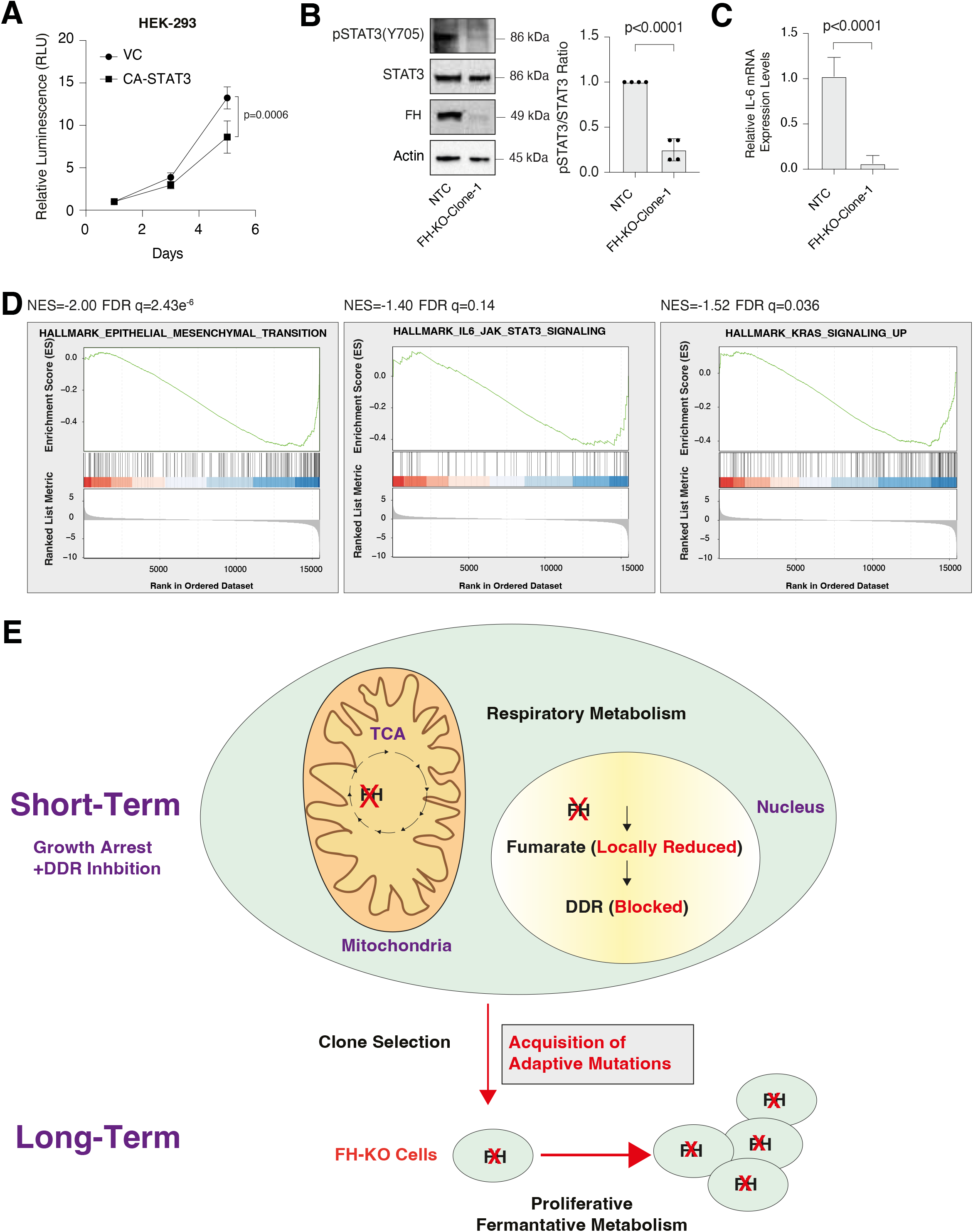
FH loss inhibits STAT3 signaling: (**A**) Constitutively activated STAT3 (CA-STAT3) inhibits cell proliferation in HEK-293T. The proliferation rates of WT and CA-STAT3 cells were measured using CellTiter Glo. Each value represents the mean ± SD for n=6. The p-value was determined by Student’s *t*-test. VC-vector control. (**B**) STAT3 phosphorylation is reduced in FH-KO-clone-1. Immunoblots representing WT HEK-293T expressing NTC and long-term FH-KO clone-1. Cells were lysed and subjected to immunoblotting using the indicated antibodies (Left). Quantification of phospho-STAT3 immunoblots normalized to total STAT3 levels; each bar represents the mean ± SD for n=3. The p-value was determined by Student’s t-test (Right). NTC -non-targeted control. (**C**) FH loss reduces IL-6 mRNA expression level. each bar represents the mean ± SD for n=12. The p-value was determined by Student’s t-test (Right). NTC - non-targeted control. (**D**) FH loss leads to reduced gene expression of the hallmark of EMT, Hallmark IL6 JAK signaling, and Hallmark of KRAS signaling. HEK-293 WT cells and FH-KO-clone-1 were subjected to RNA-Seq analysis. The expression ratio between all genes (∼22,000) was calculated and ranked based on the relative expression between the FH WT and KO. The samples were subjected to GSEA. GSEA computed FDR q-value. (**E**) A scheme demonstrating the cellular behavior at the early stages of FH knockout and its long-term result in selected clones. Immediately after FH KO, the cell proliferation rate decreases as the cells lose their ability to utilize the TCA cycle and to activate the DNA damage response (DDR) (Short-Term). This blockage in the DDR induces mutagenesis that in turn results in the acquisition of adaptive mutations. By accumulating these mutations, the cell regains its proliferative abilities.

## Discussion

The FH enzyme is a critical component of the TCA cycle and thus plays a vital role in cell metabolism and proliferation. However, cells are capable of proliferating without this enzyme, and moreover, mutation or deletion of FH is an apparent driver of various cancers ^18^. To illuminate this apparent paradox, we show that the immediate impact of FH loss is the inhibition of cell proliferation. Yet, stable knockout clones that we generated, display insignificant differences in their proliferation rate compared to the WT cells. In this study, we set out to resolve this paradox and reveal the molecular mechanism by which cells shift from FH-dependent to FH-independent proliferation.

Addressing this paradox, we describe a comprehensive model that differentiates between the short-term and long-term effects of the FH knockout. We determine that the connection between these two phases of proliferation lies in the DNA damage response, which involves the moonlighting function of FH. In WT cells, FH participates in the TCA cycle, as well as in the DNA damage response ^19^. However, upon the initiation of FH loss, the halted cells also become deficient in their ability to repair their DNA and therefore are prone to accumulation of mutations (Figure 5E). We found that these are not random mutations, as they are enriched in genes encoding signaling molecules. The STAT3 pathway is an example of a signaling cascade that is impaired in these long-term FH stable clones.

Over the past twenty years, several mechanisms have been proposed to address the FH role in cancer, mainly focusing on fumarate accumulation due to the enzyme loss ^3,20^. In renal tumors, fumarate accumulates and functions as an oncometabolite ^21^, which stimulates cancer-associated processes. For example, fumarate accumulation inhibits PH (prolyl-hydroxylase) activity and thus stabilizes HIF1α at normal oxygen tensions ^6,22^. Additionally, under acidic conditions, fumarate interacts with cysteine thiol groups, forming a post-translation modification, which regulates the activity of pro-oncogenic signaling factors such as KEAP1 (Kelch Like ECH Associated Protein 1) ^18^. We found that fumarate intracellular levels did not change significantly at the early stage of FH loss and the addition of mono-ethyl fumarate esters does not alleviate short-term FH KO effects. This indicates that the proliferation inhibition is not due to vast intracellular oncometabolite accumulation. Previously we demonstrated that FH is also found in the nucleus, where it functions in the DDR ^1,9^. Specifically, FH is localized to the damage site where it produces fumarate from malate ^4,10^. However, it is becoming clear that fumarate may have both pro and anti-DNA damage repair roles ^11,23^. We do not wish to dwell on this question regarding fumarate function in the DDR, as long as its role in this process is established.

The execution of the EMT program induces significant changes in the cell’s characteristics, such as their morphology and migratory capabilities ^24^. Notably, the EMT program outcome is reversible, as the cells can regain their epithelial characteristics by inducing the mesenchymal-epithelial transition program (MET) ^25^. Previously Sciacovelli et al. demonstrated that FH loss in mouse *Fh1*^*-/-*^ and human FH-deficiency cells (UOK262) induced the EMT program ^7^. However, our study found that in HEK-293T cells, FH loss induces the opposite MET program. Additionally, we demonstrated that FH loss affects the STAT3/IL6 signaling pathway, a central regulator of the EMT program ^26^. This inconsistency in the FH role in the cell fate could be due to the different cell line models used. Specifically, we investigated the effect of FH loss in the non-cancerous cell line (HEK-293T), which is in a hybrid state as it expresses both epithelial and mesenchymal markers ^27^. In contrast, the cells used in the other studies examined FH loss in epithelial cells. Thus, it is possible that the FH role in cell plasticity is not limited to the EMT program but has a more complex function.

## Materials and methods

### Cell Lines and Cell Culture

The cell lines HEK-293T and HepG2 were procured from ATCC and were cultured in DMEM supplemented with 10% FBS (Biological industries). All cells were cultured at 37°C with 5% CO_2_.

### Cell Lysis and Immunoblotting

Cells were rinsed twice with ice-cold PBS and lysed with RIPA lysis buffer (20 mM Tris pH 7.4, 137 mM NaCl, 10% glycerol, 1% Triton X-100, 0.5% deoxycholate, 0.1% SDS, 2.0 mM ethylenediaminetetraacetic acid (EDTA, pH 8.0), EDTA-free protease inhibitor (1 tablet per 50 ml RIPA) (Roche) and phosphatase inhibitor cocktail (100X) (Bimake). The lysates were cleared by centrifugation at 13,000 rpm at 4°C in a microcentrifuge for 10 minutes. The protein concentration was determined by Bradford (BioRad). Proteins were denatured by adding SDS sample buffer (5X) and boiling for 5 minutes, then resolved by 10% SDS-PAGE, transferred onto a 0.45 µm PVDF membrane (Merck), and probed with the appropriate antibodies.

### Exome Sequencing preparation and analysis

The DNA was extracted from HEK-293T cells using the DNeasy Blood & Tissue Kit from QIAGEN (#69504). The Samples were prepared and sequenced in Bio Basic Inc, (Singapore). The whole-exome sequencing results were analyzed by our faculty bioinformatics unit. Mutect2 (from the GATK tools) was used for calling somatic short mutations (single nucleotide (SNA) and insertions/deletions). Mutect2 uses a Bayesian somatic genotyping model and uses the assembly-based machinery of GATK’s HaplotypeCaller. We used the “Tumor with matched normal” mode with a joint analysis of multiple samples. Samples of FH KO clones were all defined as tumor samples while the WT cell line was defined as normal. The resulting vcf was further filtered using the FilterMutectCalls tool with default parameters. Variants that passed the filter were annotated by wANNOVAR online annotation tool ^14^. Only variants that were found to be located within exonic regions were kept.

### Antibodies

Antibodies were obtained from the following sources: FH and Phospho-Histone H2A.X-Ser139 (20E3 (#9718), p-sSTAT3-Tyr705 (#9145), STAT3 (#9139), Rabbit β-actin (#4970), mouse β-actin (3700) from Cell signaling Technology, and HRP-labeled anti-mouse (Jackson ImmunoResearch Laboratories, 115-035-003) and anti-rabbit (111-035-144) secondary antibodies from Jackson ImmunoResearch. For immunofluorescence assays: Goat-anti mouse-Alexa fluor 647 (Abcam, ab150119), Donkey-anti rabbit-Rhodamine red X (Jackson ImmunoResearch Laboratories, 711-295-152), and Donkey anti-rabbit-Alexa fluor 488 (Jackson ImmunoResearch Laboratories, 711-545-152).

### Virus Production

HEK-293T cells were co-transfected with the pcw107-V5 plasmid expressing CA-STAT3 (Addgene #64611), VSV-G envelope plasmid, and Δvpr lentiviral plasmid using X-TremeGene 9 Transfection Reagent. The supernatant containing the virus was collected 48 hours after transfection and spun for 5 min at 400g to eliminate cells.

### Fluorescence Microscopy

The HEK-293T (60,000 cells) and HepG2 (50,000 cells) were seeded on polylysine-coated glass coverslips in 12-well tissue culture plates. After 24 hours, the slides were rinsed twice with PBS and fixed with 4% paraformaldehyde in PBS for 15 min at room temperature, followed by quenching with ammonium chloride (1% in PBS). The slides were then rinsed three times with PBS, and cells were permeabilized with 0.1% Triton X-100 in PBS (PBS-T) for 10 minutes. The cells were gently rinsed three times with PBS and subsequently incubated for 1 hour in the blocking buffer (1% BSA in PBS-T), followed by primary antibodies for 2 hours (FH and γH2AX 1:200 each in 1%BSA in TBST) at room temperature. The cells were rinsed three times with PBS, incubated with secondary antibodies (diluted 1:200 in 1% BSA in TBST), and Phalloidin-iFluor 555 Reagent (Abcam 1:5000) for 1 hour at room temperature in the dark, and washed three times with PBS. Slides were mounted on glass coverslips using Vectashield (Vector Laboratories, # H-1000). The cells were imaged on Nikon Spinning Disk/ high content screening system using the 60X (NA=0.95, dry, CFI Plan-Apochromat Lambda) and 100X (NA=1.4, oil, Plan-Apochromat). This microscope is equipped with a spinning disk Yokogawa W1 Spinning Disk, 2 SCMOS ZYLA cameras, and 405 nm, 488 nm, 561nm, and 638nm laser. While the 20X (NA=0.5, dry, WD 2.1mm, pH1) images were acquired using the Eclipse NI-U upright microscope (Nikon), equipped with DS-QI2 MONO cooled digital microscope camera 16MP. The image analyses were done using the NIS Elements software package for multi-dimensional experiments and exported as 16-bit. The pictures were slightly adjusted (levels) using Adobe Photoshop.

### CRISPR/Cas9-Mediated Knockout Cell lines

We used CRISPR-Cas9-mediated genome editing to achieve gene knockout, using pSpCas9(BB)-2A-Puro (PX459) V2.0 (Addgene Plasmid #62988) and PX458-GFP (Addgene Plasmid #48138). Transfected cells for the long-term experiments were then subjected to single-cell cloning by limiting dilution in 96-well plates. Editing of the FH locus was confirmed by assessing protein levels by Western Blot. Primers used for cloning non-targeting control (NTC), NTC-sgRNA 5’ caccgGCGCTTCCGCGGCCCGTTCAA 3’ and 5’ aaacTTGAACGGGCCGCGGAAGCGg; 3’; FH-gRNA1 5’ caccgACATGATCGTTGGGATGCAC 3’ and 5’ aaacGTGCATCCCAACGATCATGTc 3’; FH-gRNA2 5’ caccgGGTATCATATTCTATCCGGA 3’ and 5’ aaacTCCGGATAGAATATGATACCc 3’; FHgRNA3 5’ caccgAATAATGAAGGCAGCAGATG 3’ and 5’ aaacCATCTGCTGCCTTCATTATTc 3’; DPYD-gRNA1 5’ caccgTGTGCTCAGTAAGGACTCGG 3’ and 5’ aaacCCGAGTCCTTACTGAGCACAc 3′/

### Cell Proliferation Assay

For the IncuCyte live-cell imaging system (the cells were seeded in 96-well plates at a density of 800 cells/well (HEK-293T or HepG2, WT, FH-KO short time, and stable clones). The cell proliferation rate was monitored every two hours at intervals using the IncuCyte live-cell imaging system (Sartorius, Germany). After 48 hours for HEK-293T or 70 hours for HepG2, the plates were removed and the proliferation assay results were compiled from 6 wells from each group and data were analyzed and plotted as a graph. For Cell Titer-Glo the cells were seeded in white 96-well plates (Greiner) at a density of 800 cells/well. Cell viability was assessed with Cell Titer-Glo Cell Titer-Glo (Promega, # G7571) after 1-, 3-, and 5-days following seeding, and luminescence was measured with Cytation 3 Multi-Mode Reader (Biotek).

### RNA Preparation, qPCR Analysis, and RNA-seq

Total RNA was isolated from cells using the NucleoSpin® RNA Kit (MACHEREY-NAGEL, Germany), and reverse-transcription was performed using qScript cDNA Synthesis Kit (Quantbio, USA). The resulting cDNA was diluted in DNase-free water (1:10) before quantification by real-time quantitative PCR. The mRNA transcription levels were measured using 2x qPCRBIO SyGreen. Blue Mix Hi-ROX (PCR Biosystems) and StepOnePlus (Applied Biosystems). The data expressed as the ratio between the expression level of IL-6 mRNA and that of Actin. The list of primers used for the qPCR analysis was obtained from Integrated DNA Technology: IL-6 Forward-ACTCACCTCTTCAGAACGAATTG, Reverse-CCATCTTTGGAAGGTTCAGGTTG; β-Actin Forward-CCAACCGCGAGAAGATGA, Reverse-CCAGAGGCGTACAGGGATAG. For RNA-seq analysis the RNA quality was determined using a Tap station. Only RNA samples with 9-10 RIN were subjected to further RNA-seq analysis at our faculty Genomic Applications Laboratory of the Core Research Facility. The RNA-seq results were analyzed by our faculty bioinformatics unit.

### Seahorse Assay

The sensor cartridge plate was hydrated with sterile distilled water overnight at 37°C in a CO_2_-free incubator and was replaced with a calibrant solution (Agilent) for 1 hour. Cells (2×10^4^/well) were seeded in XF96 well plates (Agilent), adhered overnight, washed, and incubated with seahorse DMEM media containing 10 mM glucose, 1 mM sodium pyruvate, and 2 mM glutamine for 40 min. The cells were sequentially treated with 1 μM oligomycin (Cayman, 13995), 2 μM carbonyl cyanide-4 (trifluoromethoxy) phenylhydrazone (FCCP) (Cayman 15218), and a mixture of 1 μM antimycin A (Santa Cruz Biotechnology, sc-202467), and 1 μM rotenone (Cayman, 13995). The oxygen consumption rate (OCR) was measured using the Agilent Seahorse XFe96 Analyzer according to the manufacturer’s instructions. The oxygen consumption rate value was normalized to cell numbers per well using Agilent cytation 5 (Biotek).

### Metabolite Supplementation

FH was knocked out from Hek-293T cells and after 48 hours 1,000 cells/well were seeded in white 96 well plates for additional 24 hours. Following the cells were treated with DMSO (control), 250 µM diethyl malate (Sigma Aldrich, #W237418), or 250 µM mono-ethyl fumarate (Sigma Aldrich, #12842225G). The metabolites were added twice a day for a total of three days. Cell viability was measured using the Cell Titer-Glo (Promega, # G7571) kit, and luminescence was measured with Cytation 3 Multi-Mode Reader (Biotek).

### Comet Assay

The assay was conducted according to the Abcam comet assay kit (#ab238544) protocol. Briefly, 35 μl of suspended cells (1×10^6^/ml) was mixed with 150 μl agarose, and 35 µl of this mixture was added to each agar precoated slide. After incubation, the slides were immersed in pre-chilled lysis buffer for one hour and replaced with a pre-chilled alkaline solution (300 mM NaOH, 1 mM EDTA-Na_2_, pH 13) for 30 min, all at 4°C and in the dark. Then the slides were electrophoresed in TBE buffer for 20 min at 35 volts (1 V/cm). After the run, the slides were dried, immersed twice with pre-chilled DI-H_2_O for 2 min, dehydrated in cold 70% ethanol for 5 min, stained with vista green DNA Dye (from the kit) at room temperature for 15 min, and analyzed using a fluorescence microscope (FITC filter) (Nikon, Eclipse Ni). Quantification was performed using the software Comet Assay IV to determine the average comet tail lengths.

### Statistical Analysis

Data are shown as mean ± SD from at least three independent biological experiments. All the statistical analyses were performed using the R (version 4.2.0) or GraphPad Prism (version 8.0) statistical analysis programs. The p*-*values in most of the figures measured between the indicated samples were quantified using the unpaired two-tailed Student’s *t*-test. Data distribution was assumed to be normal but this was not formally tested. The significance of the mean comparison is present in each figure. The pie chart was generated using R package ggplot2 and the p-value was calculated using Fischer’s exact test. Chromosome with mutation location was generated using R package RIdeogram and the gene list was taken from Piovesan St. al. ^28^. One sample proportion test was applied to calculate the significance of the mutation number for each chromosome. Benjamini-Hochberg procedure was used to normalize the result.

## Figure legend

***Figure S1: FH loss inhibits STAT3 signaling:***

(**A**) DPYD was silenced in HEK-293T using the CRISPR-Cas9 system and then subjected to 24 hours of puromycin selection. At the indicated times post-selection (short-term), the cells were lysed and subjected to immunoblotting using the indicated antibodies. NTC is a non-targeted control. (**B**) Representative pictures of the proliferation assay as taken by the IncuCyte. Images representing the number of cells during 0 and 48 hours in HEK-293T and 0 and 70 hours in HepG2. Bar=400µm.

***Figure S2: FH loss inhibits STAT3 signaling:***

(**A**) Western blot indicating FH level in the 11 different FH-KO long-term stable clones. The cells were lysed and subjected to immunoblotting using the indicated antibodies. NTC is a non-targeted control. (**B**) The cumulative distribution plot of the percent of counts indicates an alternative allele in the HEK-293T cell line. (**C**) The synonymous mutation distribution pattern is represented in the different chromosomes. The mutation that occurred in more than 4 clones, is considered a frequent mutation and marked as a red triangle.

***Figure S3: FH loss inhibits STAT3 signaling:***

(**A**) Genes encoding for signaling molecules are enriched with mutations caused by FH knockdown. All of the signaling genes were defined based on the KEGG pathway database (https://www.genome.jp/kegg/pathway.html). The number of mutations (excluding synonymous) in the signaling and non-signaling genes were determined. A pie chart representing the proportion of mutations in the signaling genes versus the remaining genes. P-value is calculated by Fisher’s exact test using R. (**B**) Same analysis as A but for synonymous mutations.

## Acknowledgments

We thank the members of the YDS and OP lab. This work was supported by the Israel Science Foundation (Grant No 299/21) and the Israeli Cancer Association (Grant 20210079, Gedalia Doron funds). The Genomic Applications Laboratory of the Core Research Facility, The Faculty of Medicine, The Hebrew University of Jerusalem, Israel, performed the RNA-Seq data analysis.

