## Supplementary figures and images for "Depletion of FH, an essential TCA cycle enzyme, drives proliferation in a two-step model"

### Supplement Figure 1

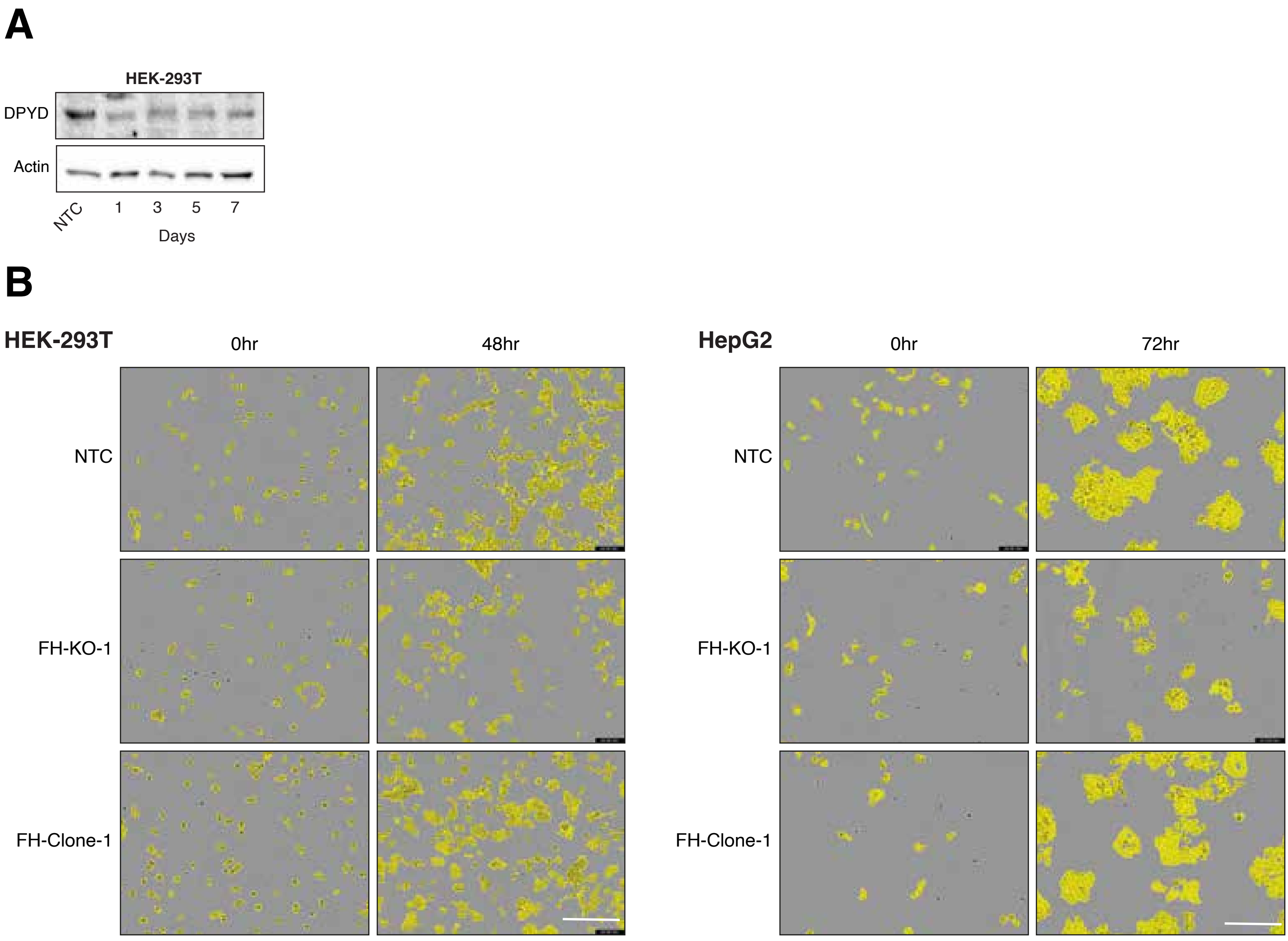

### Supplement Figure 2

A

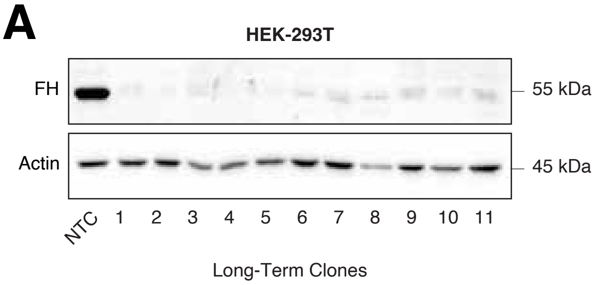


B

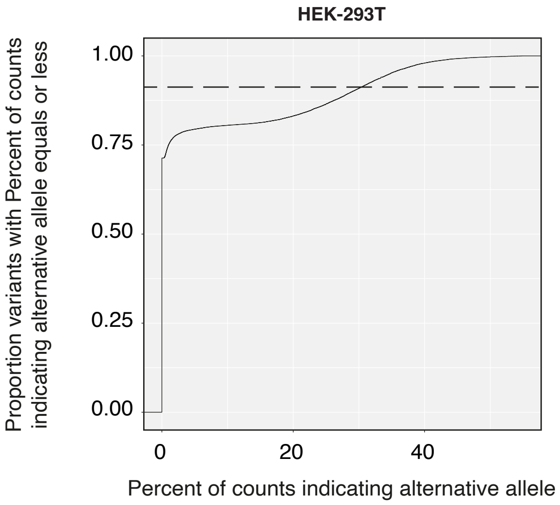


C

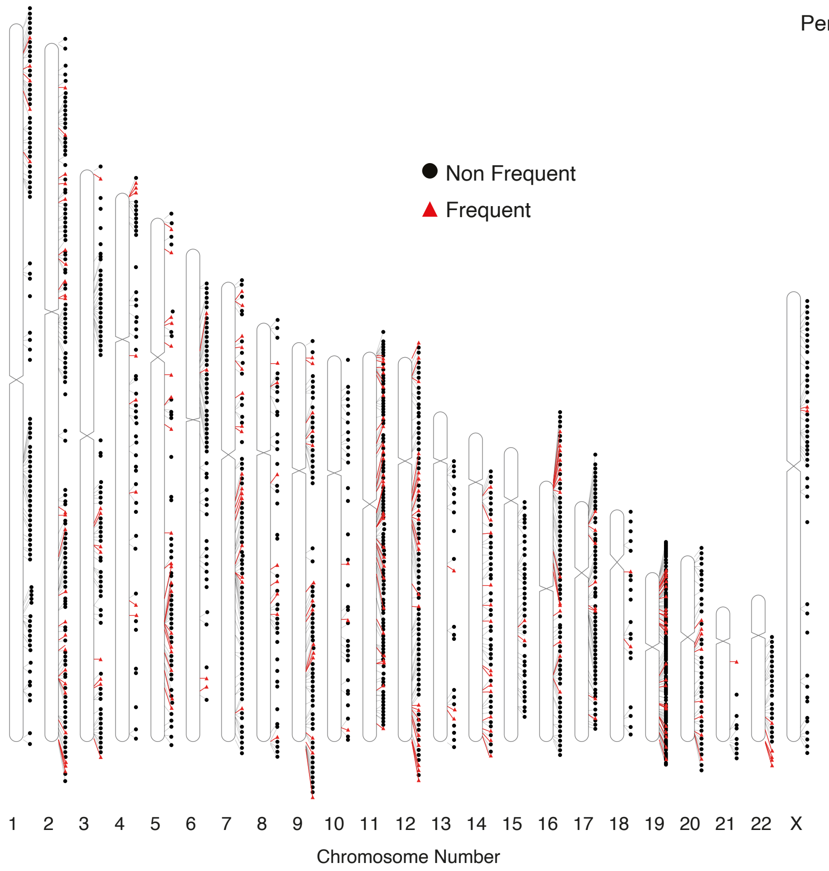


#### Figure S2

### Supplement Figure 3

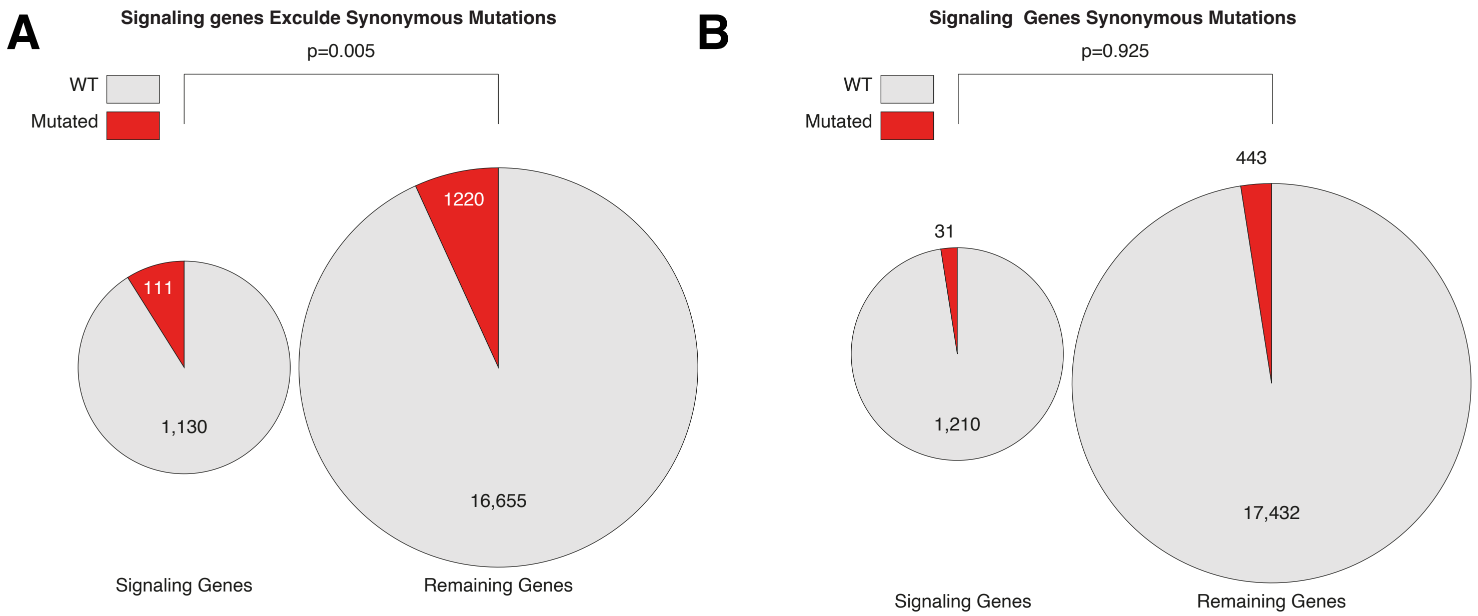
